# Sex- and estrous-specific effects of paradoxical (REM) sleep deprivation: Neurobehavioral changes and hippocampal neuroinflammation

**DOI:** 10.1101/2025.04.11.648374

**Authors:** Laura K. Olsen, Krysten A. Jones, Raquel J. Moore, Hunter McCubbins, Frances S. Curtner, Birendra Sharma, Candice N. Hatcher-Solis

## Abstract

With millions suffering from sleep disorders in today’s modern society, a better understanding of sleep disruption related cognitive outcomes is urgently needed. To that end, a preclinical investigation into the effects of sleep deprivation (SD) on neurobehavioral outcomes and associated hippocampal neuroinflammation was conducted in male and female rats. Due to epidemiological identification of sex differences in many aspects of sleep disorders, sex and estrous cycle stage factors were investigated. Sprague-Dawley rats underwent 120 h of paradoxical SD using a modified multiple platform ‘flower pot’ method. At 96 h SD, animals were trained on neurobehavioral Novel Object Recognition (NOR) and Passive-Avoidance Task (PAT) paradigms. Prior to NOR/PAT testing at 120 h SD, Elevated Zero Maze (EZM) assessed anxiolytic-like behavior. SD impaired PAT performance among males and females. Anxiolytic-like and locomotor behavior was increased and NOR performance impaired in males after SD. Estrous cycle stage determined by cytological analysis of daily wet smears found females to exhibit estrous-specific differences across all neurobehavioral paradigms, with increased anxiolytic-like behavior and impaired PAT performance only among SD females in estrus. Immunohistochemical analysis of the hippocampus after 120 h SD found microgliosis, but not astrogliosis, in the CA1/2 of males and females in estrus. This study contributes to a better understanding of sex- and estrous-specific difference in sleep disruption induced neurobehavioral outcomes and associated hippocampal inflammation. Further research is needed to investigate the molecular mechanisms underlying the interaction between estrous cycle, hippocampal microgliosis, and sleep disruption cognitive outcomes.

## Introduction

With over 50% of the U.S. population reporting sleep issues at some point in their life and 40 million suffering chronically, sleep disorders have become an urgent concern in today’s modern society [1,2]. Although sleep requirements change throughout life, both the quality and quantity of sleep are important for health and cognitive function at all ages. Nearly all age groups in the U.S. have increased reporting of sleep disorders. Roughly 60-75% of adolescents and 35% of adults report chronic insufficient sleep [3,4]. Insufficient sleep is associated with overall cognitive impairment, which can increase the likelihood of accidents and injuries while operating machinery or motor vehicles, as well as medical errors and loss of work productivity [5,6]. Disrupted sleep quality/quantity has also been associated with multiple health conditions, including metabolic disorders, cardiovascular conditions, and neurological disorders [7–10]. Although the literature suggests there are sex differences in prevalence of sleep disorders and susceptibility to subsequent disease risk associated with sleep disruption, inconsistent findings based on specific sleep dimension factors indicate that further research is needed to better understand the influence of sex on sleep disruption related cognitive outcomes [11–13].

The hippocampus mediates declarative learning and memory integration, encoding, and consolidation [14,15]. Activation of neural processes within the hippocampus lead to molecular and structural synaptic plasticity changes that underlie these cognitive phenomena. Although the behavioral significance is not fully understood, sex-specific differences in dendritic spine densities and structures of pyramidal neurons of the hippocampus suggest that sex-specific differences in synaptic plasticity may exist [16,17]. As regulators of the brain’s microenvironment, glial cells within the hippocampus modulate neuronal synaptic plasticity [18,19]. Microglia, the resident immune cells of the brain, can participate in healthy synaptic regulation or runaway pro-inflammatory response depending on their activation state [19,20]. Astroglia crosstalk with microglia during inflammatory insults also mediate the progression of neuroinflammation within the brain. Activated microglia and astroglia release pro-inflammatory mediators that modulate synaptic protein levels and dendritic spine densities [21]. Within the hippocampus, these glial cells influence learning and memory associated activity-dependent synaptic plasticity pathways [22]. Increases in activated microglia and pro-inflammatory cytokines have been found after chronic (SD) [23]. Although the current understanding from the literature identifies the pro-inflammatory glial response in the hippocampus to in part mediate cognitive impairment during disrupted sleep, the effect of sex has not yet been investigated.

Firstly, this study sought to characterize the cognitive behavioral outcomes of SD across a battery of neurobehavioral tests in male and female rodents. Female animals have historically been generally excluded from neurobehavioral investigations due to the ‘confounding’ variable of the estrous cycle [24]. In this study, the estrous cycle of the females was tracked to determine the influence of female sex hormones on cognitive behavioral outcomes of SD. Lastly, this study investigated glial cell expression in hippocampal subregions after SD in male and female rodents to determine if there are sex- or estrous-specific effects.

## Methods

The views and opinions presented herein are those of the author(s) and do not necessarily represent the views of the DoD or its Components. Appearance of, or reference to, any commercial products or services does not constitute DoD endorsement of those products or services. The appearance of external hyperlinks does not constitute DoD endorsement of the linked websites, or the information, products, or services therein.

### Animals

All animal activities were conducted in an AAALAC International accredited facility in compliance with all federal regulations governing the protection of research and animals and DODI 3216.01. This study was approved by the Wright-Patterson Air Force Base IACUC. Male or female Sprague-Dawley rats (Charles River) were group housed by biological sex with *ad libitum* access to water and food on a 12 h light cycle. Rats were acclimated to the facility for at least one week before the study began and were assigned to the control or SD groups. Rats underwent a total of 120 h of SD during behavioral testing (Figure S1A). At the time of behavior, all rats were aged 10-12 weeks. Estrous cycle was tracked in female rats by wet smear collection for at least five days during experimentation.

### Paradoxical SD

A modified version of the “flower pot” method using multiple platforms in a large container was utilized to allow animals to undergo paradoxical SD in a group housing environment and prevent stress induced by immobilization on a single platform [25]. Animals were placed on small circular platforms (5 cm in diameter) that were spaced 8 cm apart and ∼3 cm above the water level. In this condition, animals are able to rest on the small platform but if they have sleep related muscle tone loss, they are awakened by falling into the water below. To prevent hypothermia stress, the water was warmed to 26-28 °C and the ambient room temperature was ∼80 °C. Due to the reliance on loss of muscle tone to enforce wakefulness, this method is considered a model of paradoxical or REM SD [27]. Control animals were placed on large square platforms (14 cm x 14 cm) that allowed for normal sleeping behavior above the water. Food and water were provided *ad libitum*. The water containers were cleaned daily, during which time animals were placed in dry cages for 30-60 min and kept awake with grooming and exploratory behavior.

### Wet Smears

Estrous cycle stage of the female Sprague-Dawley rats was tracked by analysis of wet smear collections. Vaginal lavage was collected three times with a pipette and the same 10 µl of sterile saline. Samples were air dried on slides and hematoxylin and eosin staining was performed using a Tissue-Tek Prisma Plus Automated Slide Stainer. The slides were first immersed in hematoxylin (7017, Astral Diagnostic) and then eosin (s176, Poly Scientific R&D Corp). Next, a series of washes in denatured alcohol (HC-1100, Fisherbrand) and xylene (HC700, Fisherbrand) were completed. The stained samples were evaluated to determine cell cytology. The estrous cycle stage was classified by the ratios of leukocytes, nucleated epithelial cells, and cornified epithelial cells. Estrous cycle stage was assigned from wet smears collected on training day for cognitive neurobehavioral paradigms and immunohistochemical analyses. For analysis of the effects of estrogen, metestrus and diestrus cycle stages were grouped together as ‘diestrus phase,’ while proestrus and estrus cycle stages were grouped as ‘estrus phase.’

### Elevated Zero Maze (EZM)

The EZM test was performed to assess anxiety-like behaviors (Figure S1B). The EZM was elevated 65 cm above the floor and the circular platform (outer diameter 100 cm) was divided into four equal arms. Two opposite arms were open and the remaining two arms were closed (surrounded by 30 cm high dark walls). The open arm regions were surrounded by an edge 1 cm high. Animals were placed in the middle of a closed arm (left or right, randomized for each trial) and were allowed to explore for five minutes. During the trial, the rat’s activity was monitored using Ethovision XT (v11.5, Noldus).

### Novel Object Recognition (NOR)

The NOR paradigm was used to investigate changes in recognition memory (Figure S1C). Rats were allowed to explore the NOR arena (60 cm x 60cm x 38 cm) in the presence of two similar objects (∼8 cm radius, ∼6 cm tall crown shaped glass item spray painted black) for three min. The following day, rats were again allowed to explore the NOR arena in the presence of the same object and a novel object (∼8 cm radius, ∼10 cm tall rook shaped glass item spray painted black) for three min. The objects were 47 cm apart on opposite sides of the arena. Rats habituated to the arena for two min before the object exposure on training and testing day. Ethovision XT was used to monitor object exploration activity. Novel object preference (NP) was calculated on testing day using the following equation:

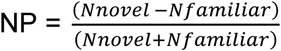

### Passive Avoidance Task (PAT)

The PAT was used to detect changes in associative aversion learning (Figure S1D). On habituation day, rats were allowed to explore the lit chamber and dark chamber (25 cm x 21 cm x 17 cm) of the apparatus (Gemini Avoidance System, San Diego Instruments Inc.) for five min. After ∼24 h, the rats were placed in the lit chamber and again allowed to enter to the dark chamber. If the rat chose to cross to the dark chamber, the gate separating the chambers closed and a one-time foot shock (0.75 mA) as delivered in the dark chamber. After the foot shock, the rat remained in the dark chamber for an additional 30 s to facilitate aversive training. The following day, the rat was returned to the lit chamber. Latency to enter the dark chamber from the lit chamber was measured and analyzed to determine associative learning retention.

### Immunohistochemistry (IHC) Staining

After completion of behavioral paradigms on testing day, rats were euthanized under deep anesthesia by exsanguination with saline perfusion. The brain was immediately extracted and fixed in 4% paraformaldehyde (PFA) for 48 h. Fixed brain tissue was then transferred to 30% sucrose for at least three days. A sliding microtome (SM2010R, Leica) was used to collect 30 µm coronal hippocampal slices for immunostaining of ionized calcium binding adaptor molecule (Iba1) or glial fibrillary acidic protein (GFAP) protein. Tissue was blocked (PBS with 2% goat serum, 0.2% Triton X-100) at room temperature for 1 h and incubated in rabbit anti-Iba1 (1:500, Wako, 019-19741) or rabbit anti-GFAP (1:2000, Agilent, Z033429-2) overnight at 4 °C. The following day, mouse anti-NeuN (1:2500, Sigma, MAB377) was added to the primary antibody incubations for 2 h at room temperature. The tissues were then rinsed five time (PBS, 5 min/rinse) before the slices were incubated in secondary antibodies (goat anti-rabbit Alexa Fluor 488 and goat anti-mouse Alexa Fluor 594, Jackson ImmunoResearch, 111-545-144 and 115-585-146) for 1 h at room temperature. The tissues were rinsed (PBS, 5 min/rinse, five rinses) and then mounted using Fluroshield with dapi (Sigma, F6057). Fluorescent images were collected using an Olympus microscope at 20x magnification and quantified using QuPath and ImageJ. NeuN was used to identify the stratum radiatum and pyramidal layer of CA1 and CA2. Quantification of Iba1 or GFAP positive cells in each region was normalized to dapi counts. IHC quantification was performed on three technical replicates for each rat biological sample by a blinded experimenter. Representative Iba1 and GFAP images for all hippocampal regions are provided in Figures S4 and S7.

### Statistical Analysis

All data were examined for normality and homogeneity of variance. Statistical significance of the behavioral data and IHC expression data sets was determined by unpaired, two-tailed *t*-test, Mann-Whitney test, or two-way between group analysis of variance (ANOVA). Post-hoc analysis was performed by Bonferroni comparison. Data are presented as means ± the standard error of the mean (SEM). A calculated p-value of less than 0.05 was considered statistically significant. Detailed statistical analysis results are provided in Tables S1-9.

## Results

### Paradoxical SD effects on anxiety measures

The EZM paradigm was used to examine the impact of paradoxical SD on anxiety-like behaviors in male and female rats. After 120 h of SD, general locomotion was statistically greater among SD male rats compared to their control group (Figure 1A). No difference in distance traveled was observed among female rats, regardless of estrous cycle stage (Figure 1B-D). Male SD rats spent more time in the open arm, which is often thought to provoke anxiety, when compared to control rats (Figure 1E). Time in the open arm was also approaching significance between all female SD and controls (Figure 1F). When estrous cycle was examined among the female rats, a significant increase in the amount of time in the open arm was only found for SD females in estrus (Figure 1G-H).

**Figure 1.**
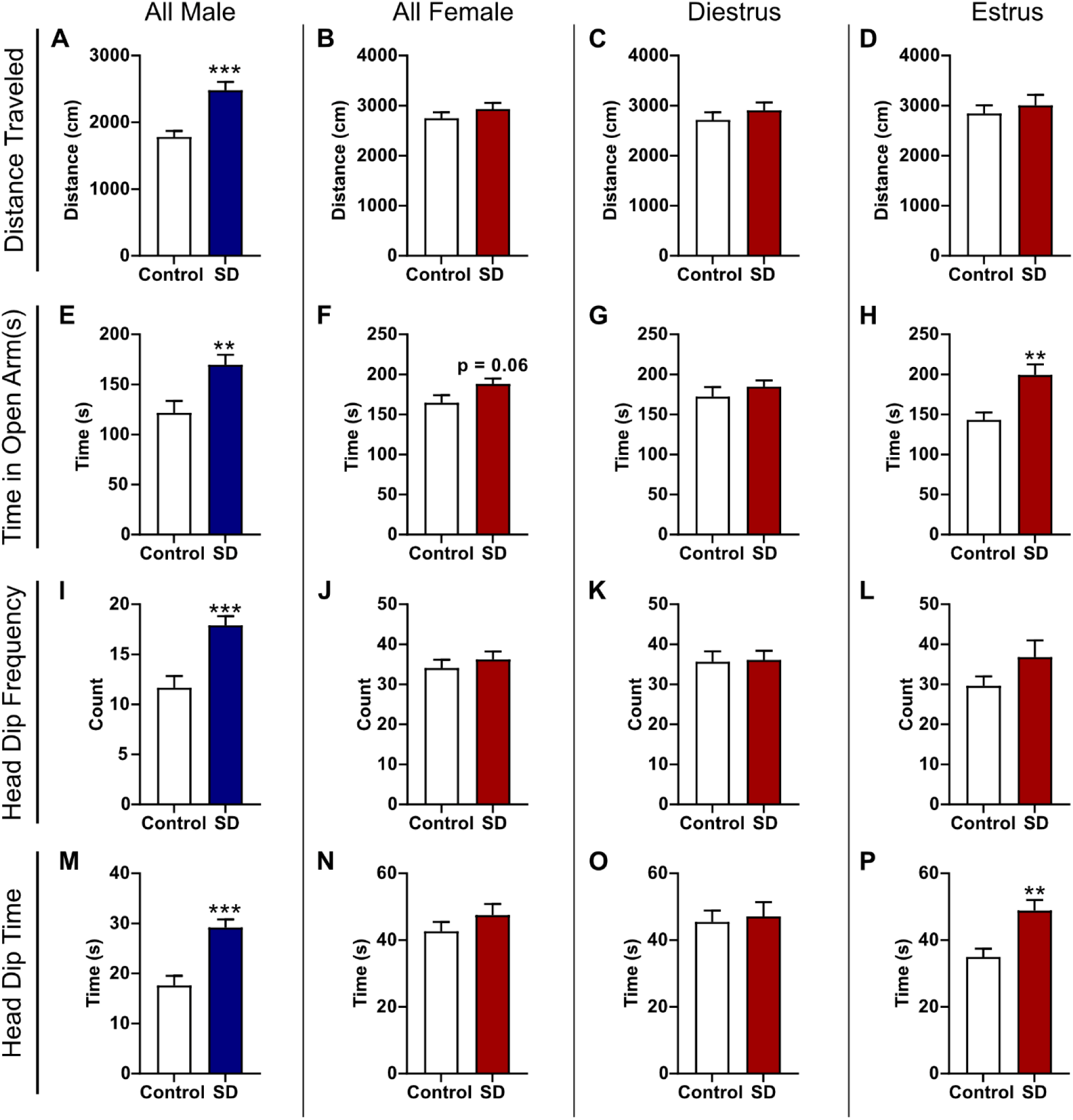
SD effects on anxiety-like behavior in the EZM paradigm are sex-dependent. (A) Distance traveled in the EZM was increased for male SD rats. (B) SD did not affect distance traveled in all female subjects or for females in (C) diestrus or (D) estrus. (E) In all male subjects, SD decreased anxiety-like behaviors as indicated by an increase in the time spent in the EZM open arm. (F) An increase in time spent in the open arm approached significance for all female SD subjects. (G) SD before EZM did not influence time spent in the open arm for females in diestrus and (H) increased time spent in the open arm for SD females in estrus. (I) Head dip frequency was increased for SD male rats. (J) Head dip frequency was not affected by SD for all female subjects, including females in (K) diestrus and (L) estrus. (M) Head dip time was increased for SD male rats. (N) Head dip time was not affected by SD for all female rats or (O) females in diestrus. (P) Head dip time was increased for SD estrus females. Bar graph data are presented as mean ± SEM. *p < 0.05 vs control, **p < 0.01 vs control, ***p < 0.001 vs control.

Head dip activity was also assessed as an additional measure of relative anxiety. Head dip frequency was significantly different between control and SD groups for all males, with the SD rats displaying more head dips than controls (Figure 1I). There was no significant difference in head dip frequency between the SD and control group among all female rats (Figure 1J). Estrous cycle stage did not significantly influence head dip frequency between control and SD females though there was a trending increase among SD rats in estrus (Figure 1K-L). Similar to head dip frequency, the total head dip time was significantly increased for SD male rats compared to the male control (Figure 1M). Head dip time was also estrous cycle dependent with only SD females in estrus at the time of testing having a significant increase as compared to the control group (Figure 1N-P). Collectively, these results indicate SD reduces anxiety-like behaviors in male and female rats. This effect is estrous cycle-dependent in female rats as significant differences in anxiety behavior are only observed in the estrus phase. However, this effect is an interaction with paradoxical SD since there was no estrous cycle-dependent effect in the control condition across all EZM measures (Figure S2A-D).

### Paradoxical SD effects on memory and learning

The NOR and PAT paradigms were used to examine the effects of paradoxical SD on recognition memory and learning/memory, respectively. After 96 h of SD, the NOR and PAT training sessions were completed. The NOR and PAT testing were completed the following day after an additional 24 h of SD. During the NOR testing session, general locomotor activity was also measured prior to introduction of the familiar and novel objects. Male SD rats had significantly greater locomotor activity than male controls (Figure 2A). No significant difference between the control and SD groups was detected among all females or based on estrous cycle phase (Figure 2B-D). These results are consistent with the level of locomotor activity observed by distance traveled in EZM.

**Figure 2.**
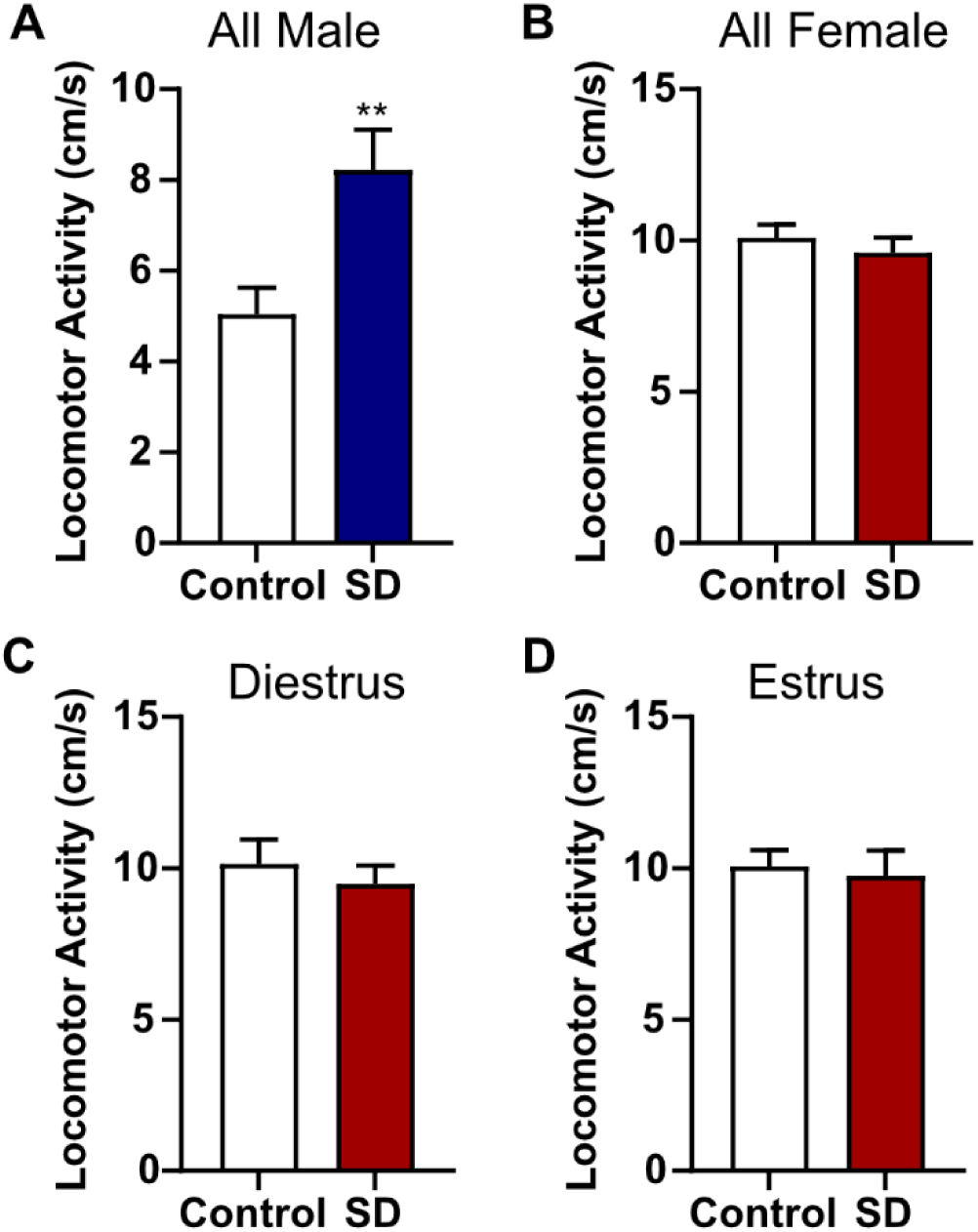
Locomotor activity is only increased by SD in male rats. (A) Locomotor activity was increased for male SD rats compared to controls. (B) SD did not affect locomotor activity in all female subjects or for females in (C) diestrus or (D) estrus. Bar graph data are presented as mean ± SEM. **p < 0.01 vs control.

When the familiar and novel objects were introduced on the NOR testing day, there was no significant difference in object exploration frequency for SD male rats (Figure 3A) or SD female rats (Figure 3B). These effects were mediated by estrous cycle stage in the female rats, as there was no difference in frequency of object exploration for SD females in diestrus but SD females in estrus more frequently explored the novel object (Figure 3C-D). For object exploration time, a two-way ANOVA found a statistically significant interaction between the effects of group and object in male rats. Post-hoc analysis revealed the male control group spent significantly more time exploring the novel object than the familiar object (Figure 3E). There was also a trending decrease in novel object exploration time for the male SD group. There was no significant difference in the time spent with the familiar or novel object among all female subjects (Figure 3F). Similar to the frequency of object exploration, the time exploring objects was also estrous cycle dependent. There was no significant difference in the time with the novel or familiar object for control and SD females in diestrus (Figure 3G). In contrast, only SD females in estrus spent significantly more time with the novel object compared to the familiar object (Figure 3H). When normalized for total exploration time, there was a significant decrease in novel object preference (NP) for the male SD group compared to the controls (Figure 3I). NP was not significantly different among the female rats regardless of estrous cycle (Figure 3J-L).

**Figure 3.**
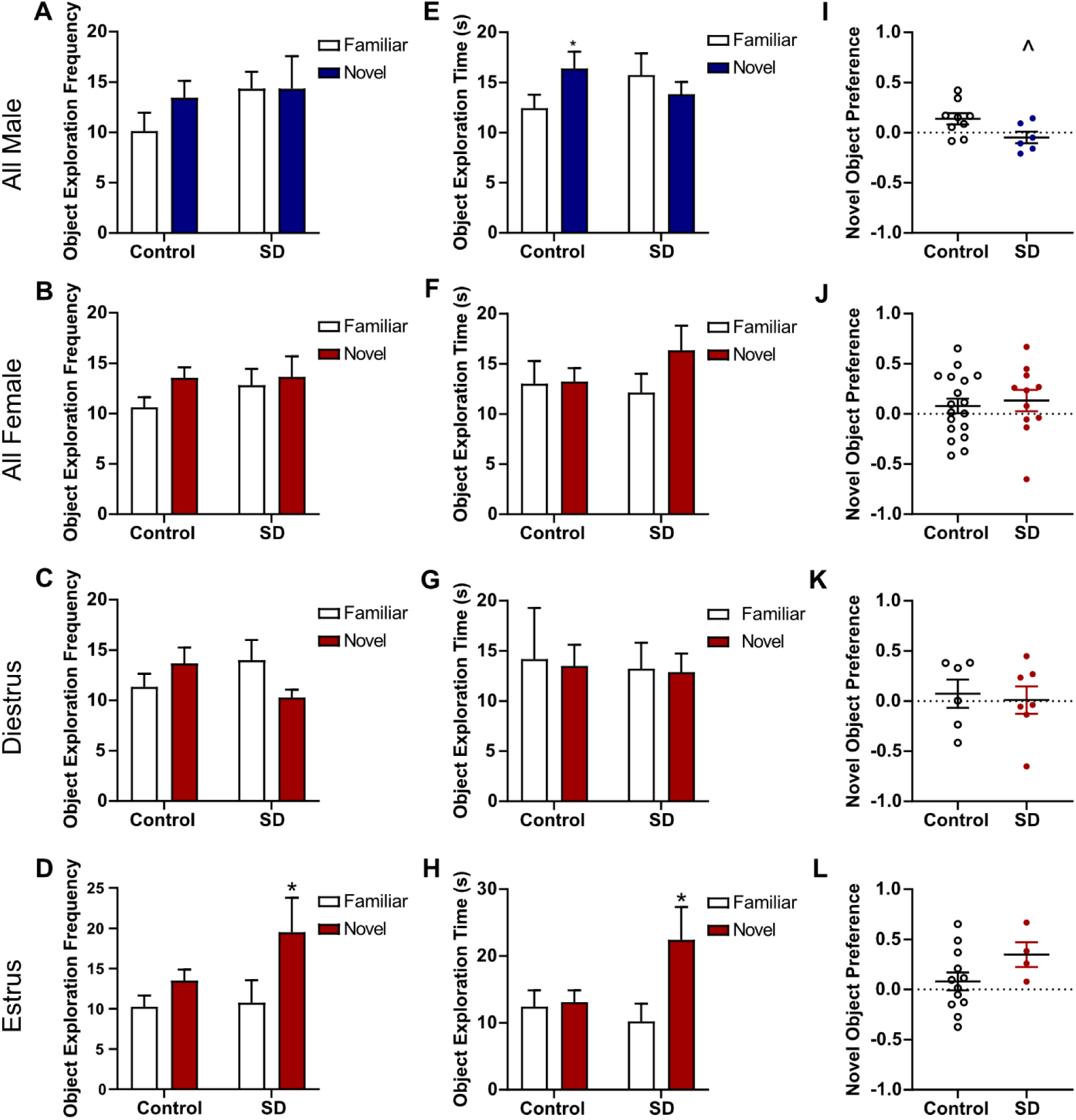
NOR performance after SD: sex and estrous cycle mediate SD effects. (A) No significant effects of SD before NOR testing were found for object exploration frequency by a two-way ANOVA for all male subjects. (B) A two-way ANOVA also found no significant effect for object exploration frequency for all female subjects and (C) females in diestrus. (D) For females in estrus, a significant main effect of ‘Object’ for frequency of object exploration was found by two-way ANOVA. Post-hoc analysis found this effect was significant for the SD estrus females. (E) A two-way ANOVA found a significant interaction between group and object for object exploration time in male subjects. Post-hoc analysis found the effect was significant among the control animals. (F) No significant effects of SD were found for object exploration time by two-way ANOVA for all female subjects and (G) females in diestrus. (H) A two-way ANOVA found a significant interaction between group and object and a significant main effect of ‘Object’ for object exploration time for females in estrus phase. Post-hoc analysis found the effect was significant among the SD animals. (I) SD before NOR testing caused a decline in NP score for male subjects. (J) SD did not affect NP for all female subjects and for females in (K) diestrus or (L) estrus phase. Bar graph and scatter plot data are presented as mean ± SEM. *p < 0.05 vs familiar object, p < 0.05 vs control.

The PAT was used to examine the effect of SD on associative aversion learning and memory. PAT performance was significant declined for male SD rats when compared to the control group (Figure 4A). Similarly, all SD females performed worse in the PAT as compared to control female rats (Figure 4B). PAT performance was also influenced by estrous cycle stage as there was no significant difference between the control and SD females in diestrus (Figure 4C). In contrast, SD females in estrus were found to have decreased PAT performance when compared to the control estrus group (Figure 4D). Although estrous cycle stage influenced NOR and PAT behavioral outcomes among sleep deprived females, comparisons between females in diestrus and estrus in the control group indicate that estrous cycle alone does not modulate behavior in these paradigms (Figure S2E-I).

**Figure 4.**
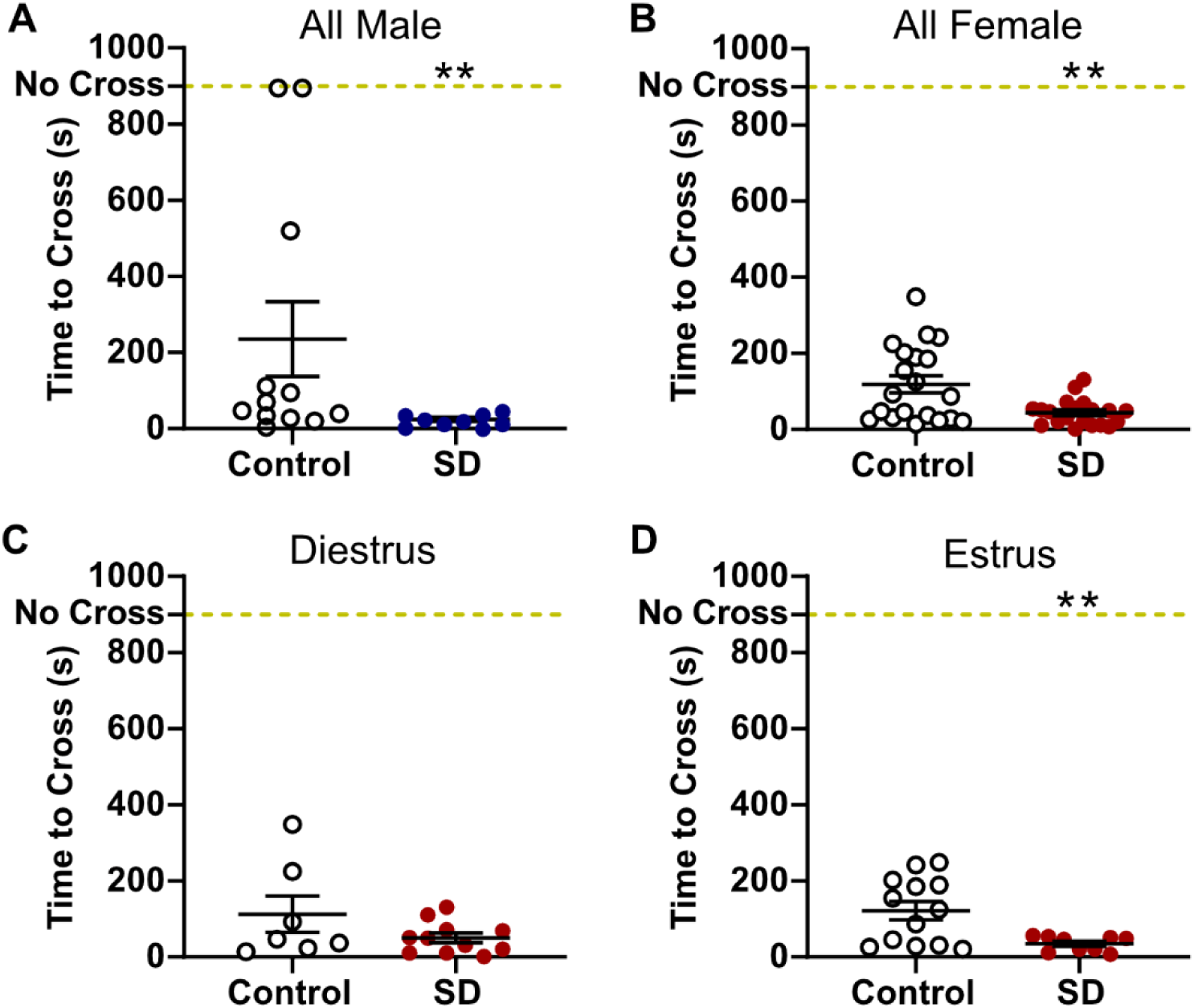
PAT performance after SD is not dependent on sex and influenced by estrous cycle. (A) SD decreased PAT performance in all male subjects. (B) PAT performance was also decreased among all female SD rats. (C) No change in PAT performance was found among females in diestrus. (D) The decline in PAT performance was maintained in female SD rats in estrus phase. Scatter plot data are presented as mean ± SEM. **p < 0.01 vs control.

### Hippocampal specific changes related to learning and memory performance

To investigate glial cell expression changes associated with decreased learning and memory performance after SD, IHC was conducted to determine changes in the microglia marker ionized calcium binding adaptor molecule 1 (Iba1) and astroglia marker glial fibrillary acidic protein (GFAP) in the hippocampus. Iba1 expression in the CA1 stratum radiatum (SR) was increased in the SD groups for both all male and all female subjects when compared to the respective control groups (Figure 5A-B). While there was a trending increase in CA1 SR Iba1 expression among SD females in diestrus, the increased expression was only significant for SD females in estrus (Figure 5C-D). SD also significantly increased Iba1 expression in the CA2 SR when compared to controls for both the male and all female subjects (Figure 5E-F). Increased Iba1 expression in the CA2 SR was also estrous cycle dependent as there was no significant difference between the control and SD diestrus groups while the increased Iba1 was maintained for SD estrus females (Figure 5G-H). Analysis of Iba1 expression in the pyramidal layer (PL) of CA1 and CA2 found no significant changes in expression between the control and SD groups for either sex or estrous cycle (Figures S3-4). Similarly, SD did not change the total GFAP expression in the SR or PL in the CA1 or CA2 based on sex or estrous cycle stage (Figures S5-7).

**Figure 5.**
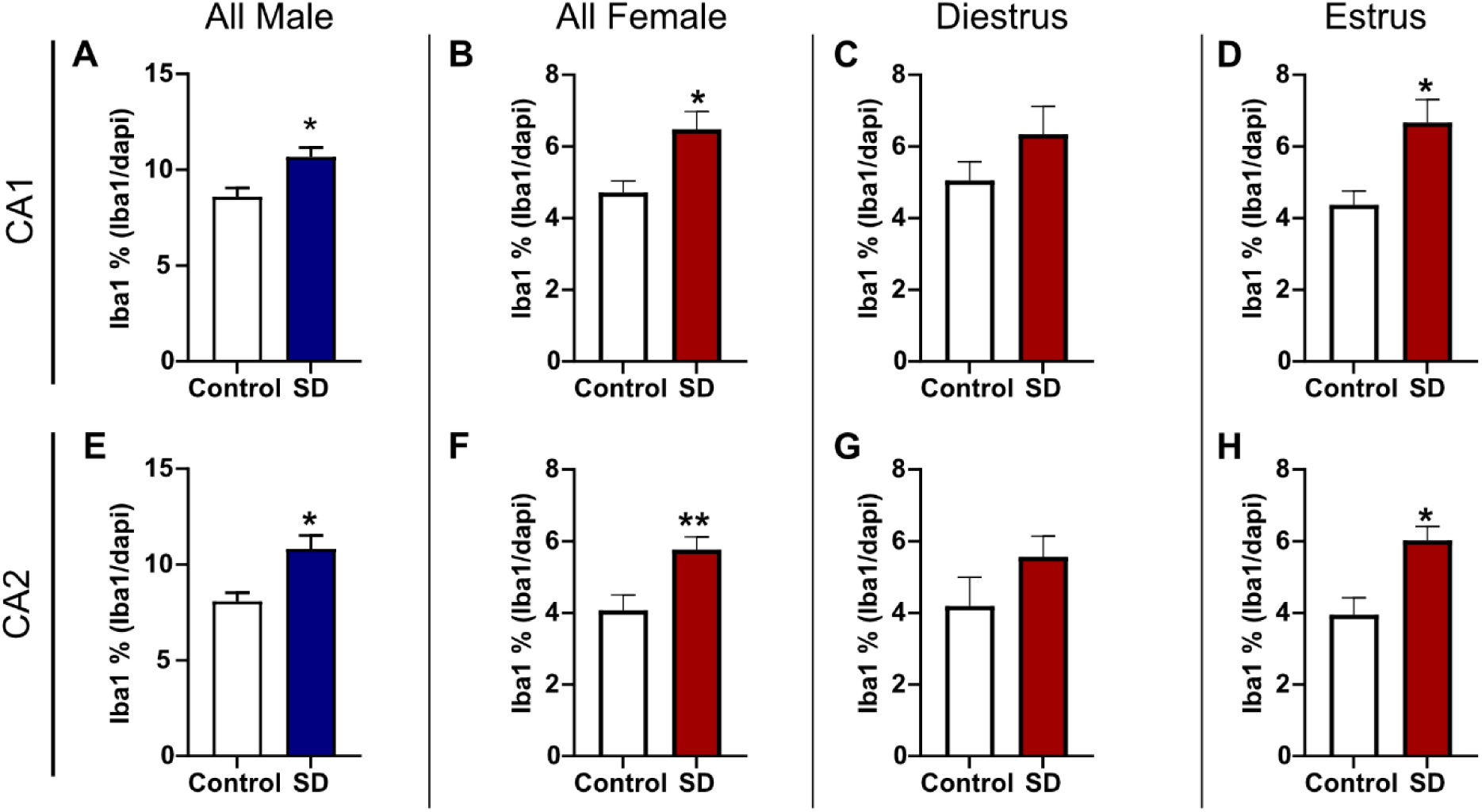
Iba1 expression in the hippocampus increases after 120 h SD. (A) Iba1 expression increased in the CA1 SR in all male rats after SD. (B) SD increased Iba1 expression in the CA1 SR in all female rats. (C) There was no change in Iba1 expression among female rats in diestrus phase while (D) the Iba1 increase in the CA1 SR was maintained for female SD rats in estrus. (E) Iba1 expression in the CA2 SR was significantly increased between control and SD male subjects. (F) SD also induced an increase in CA2 SR Iba1 among all female rats. (G) No significant difference in Iba1 expression was found in the CA2 SR for females in estrus. (H) SD increased Iba1 expression in the CA2 SR among females in estrus phase. Bar graph data are presented as mean ± SEM. *p < 0.05 vs control, **p <0.01 vs control.

## Discussion

In this study, we aimed to characterize the behavioral effects of SD in male and female rats with a focus on cognitive outcomes. Overall, disrupted sleep led to anxiolytic-like effects, impaired learning, and hippocampal microgliosis in males and females in estrus. Results suggest that although there are sex-specific differences in paradoxical SD outcomes, the primary mediator underlying these sex-specific differences are estrous cycle-dependent effects among females. An interaction between estrous cycle stage and paradoxical SD determines behavioral and neuroinflammatory outcomes among females. Although results suggest that microgliosis in the hippocampus may play a factor in the manifestation of paradoxical SD-induced behavioral effects among males and females in estrus, whether this is association is causative or symptomatic of other upstream pathways requires further investigation.

Previous animal research on anxiety-like behavior after SD is laden with inconsistent findings [27–29]. A meta-analysis concluded that preclinical models of SD decrease anxiety-like behavior [29] and the general consensus of the analysis aligns with the present study’s findings. However, they caution the translatability of traditional anxiety paradigms used in preclinical studies for the purposes of sleep research [29]. Intriguingly, a previous study found exercise to modulate the effects of SD on anxiety-like behavior [27]. The males in the present study exhibited consistent increased locomotor activity in EZM and NOR suggesting the opportunity for exercise based on the SD paradigm type may explain our results and inconsistencies in previous preclinical studies. However, an increase in anxiolytic-like behavior without increased locomotor activity among females in estrus suggests sex- and estrous-specific differences mediate sleep disruption-induced anxiety-like behavioral outcomes. Previous studies comparing sex-specific differences found females to be more susceptible to increased anxiety-like behavior after SD [28,30]. Although few studies have investigated the influence of estrous cycle on anxiety-like behavior in rodents, some studies have reported reduced anxiety-like behavior among females in proestrus/estrus [31–33]. Although this study did not replicate this finding when comparing diestrus and estrus females in the control group, this previously described phenomena might explain the interaction found between estrous cycle stage and paradoxical SD.

Previous research on the effects of SD on learning and memory are also inconsistent, especially when sex- and estrous-dependent effects are investigated [28,34,35]. The influence of sex on SD-induced cognitive impairment appears to be task dependent. Previous studies found females to be more susceptible to SD-induced cognitive performance impairment in the Morris Water Maze and discrimination tasks [28,34]. However, in alignment with this study’s findings, no sex specific difference in PAT performance after SD was found [34]. Similar to the literature, the overall cognitive effects of SD were not consistent among females between NOR and PAT. This could be due in part to the lack of preference for the novel object among control group females. A change towards impaired performance in NOR behavior cannot be detected if the control group does not demonstrate a novel object preference. This suggests that the NOR paradigm in this study might need to be modified (such as increasing the amount of exploration time during training) for this group. Although not replicated in this study, a previous study found females in estrus to perform better in a discrimination task compared to females in diestrus that was abolished by SD [35]. The present study’s finding that estrous cycle mediates the effect of paradoxical SD on fear aggravated learning in PAT hasn’t previously been investigated to our knowledge. Although previous studies implicate estrogen as neuroprotective, these studies were often conducted in ovariectomized females or disease conditions with conclusions that may not apply to the context of SD [36–38].

Glial cell expression in hippocampal subregions was investigated after paradoxical SD to determine if there are sex- or estrous-cycle dependent effects and if these effects parallel the neurobehavioral findings. Interestingly, the Iba1 results indicate that paradoxical SD-induced microgliosis patterns in the CA1/2 corresponded to multiple neurobehavioral outcomes. The same males and females in estrus that exhibited increased anxiolytic-like behavior and cognitive impairment in a fear aggravated learning task had increased microglia presence in the CA1/2. A previous study in rats also found hippocampal microgliosis and increased pro-inflammatory cytokines after SD that was associated with impaired spatial memory performance [39].

The novel contribution of the present study is the finding that paradoxical SD-induced microgliosis is only sex-specific in the context of estrous cycle-dependent effects. Previous studies indicate that microglia have estrogen receptors and that exposure to estrogen hormones modulate microglia expression of immune regulating major histocompatibility complex-I (MHC1-) [41]. Based on the literature, estrogen hormone mediated suppression of MHC-1 would prevent microglia synaptic pruning [41,42]. Although under certain circumstances preventing microglia synaptic pruning may be neuroprotective, in the context of SD and sleep associated homeostatic synaptic plasticity, a suppression of microglia synaptic pruning may hinder normal memory consolidations processes [42–44]. Further research is required to better understand the association between hippocampal microgliosis and SD neurobehavioral outcomes, especially in the context of sex- and estrous-specific differences.

Overall, the present study contributes to a better foundational understanding of sex- and estrous-specific outcomes after SD. The inclusion of female subjects and investigating estrous-cycle effects is still relatively in its infancy. In addition to contributing to the field of sleep research, this study further supports the need to re-examine neurobehavioral concepts with a focus on sex-specific and estrous-cycle investigation.

## Acknowledgements

This work was supported by the Air Force Office of Scientific Research of the United States (AFOSR grant number 20RHCOR04). The authors would like to thank the Wright-Patterson Air Force Base Research Support Command for their assistance with this work.

## Data Availability

The data underlying this article are available from the corresponding author upon reasonable request.

