## Supplemental Figures and Tables for "Sex- and estrous-specific effects of paradoxical (REM) sleep deprivation: Neurobehavioral changes and hippocampal neuroinflammation"

**Table of Contents**

| | |
|---|---|
| Figure S1 | S2 |
| Figure S2 | S3 |
| Figure S3 | S4 |
| Figure S4 | S5 |
| Figure S5 | S6 |
| Figure S6 | S7 |
| Figure S7 | S8 |
| Table S1 | S9 |
| Table S2 | S10 |
| Table S3 | S10-11 |
| Table S4 | S12 |
| Table S5 | S12 |
| Table S6 | S13 |
| Table S7 | S13 |
| Table S8 | S14 |
| Table S9 | S15 |

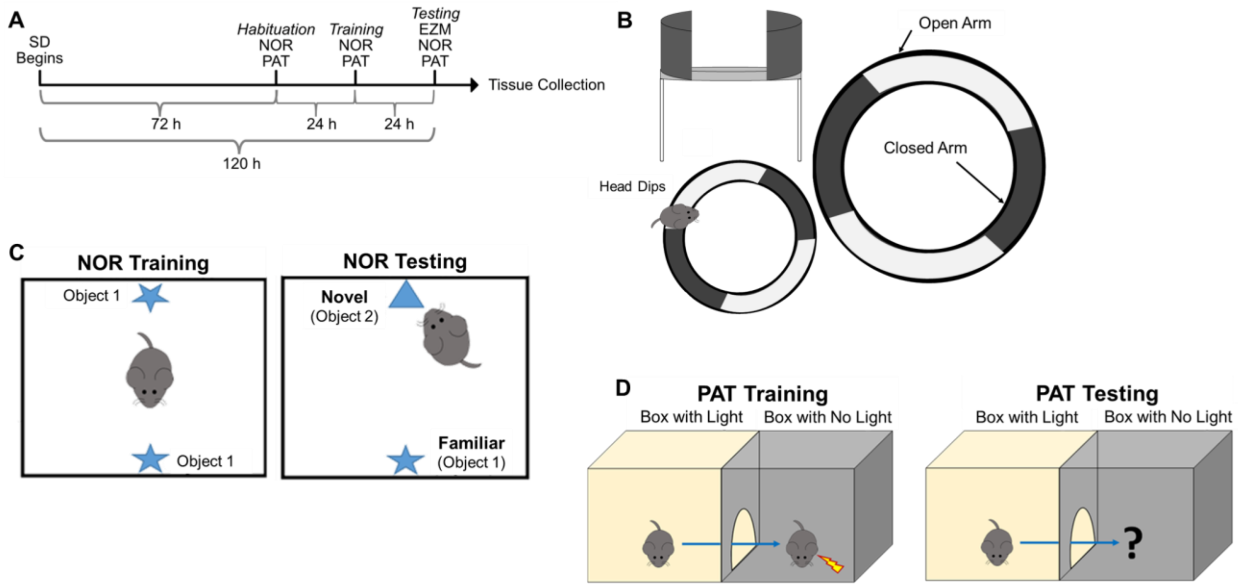


**Figure S1. Overview of experimental design.** (A) Male and female Sprague-Dawley rats underwent a total of 120 h of sleep deprivation (SD) during behavioral testing. After 72 h of SD, animals were habituated to the NOR and PAT arenas on habituation day. The following day, after an additional 24 h of SD, training for the NOR and PAT tasks was completed. Testing for the (B) EZM, (C) NOR, and (D) PAT paradigms was performed after an additional 24 h of SD. Animals were euthanized and brain tissue was collected after SD and behavioral testing for immunohistochemistry (IHC). Control animals were subjected to the same experimental timeline but control platforms were utilized that do not restrict sleep.

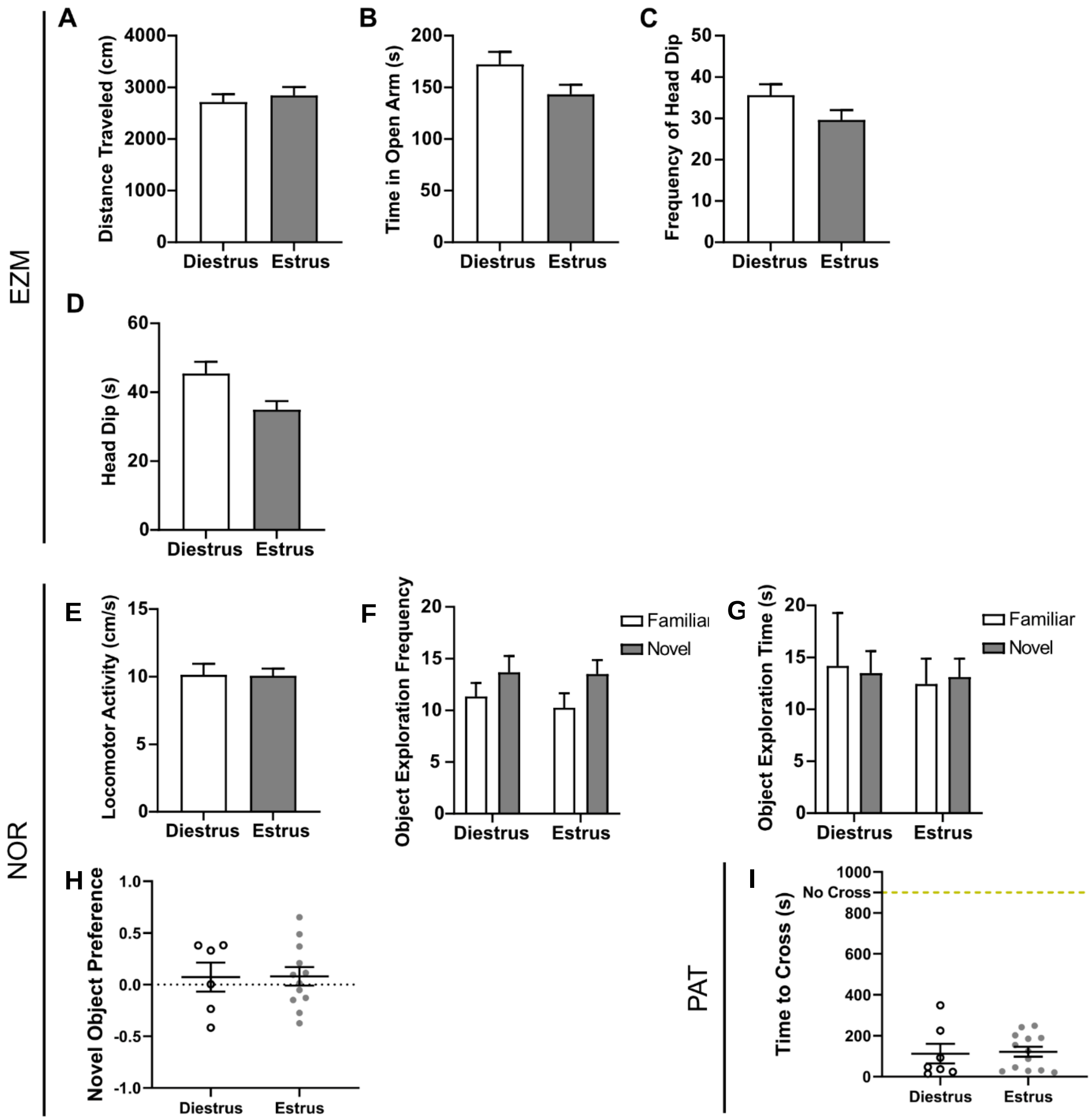


**Figure S2. Behavior outcomes are not significantly different between estrous cycle stage for control animals.** (A-D) Measurements of anxiety-like behavior as assessed by EZM did not change between control diestrus and control estrus females. (F-I) Estrous cycle did not change locomotor activity or NOR performance among female controls. (J) PAT performance as not statistically different between diestrus controls and estrus controls. Bar graph and scatter plot data are presented as mean ± SEM.

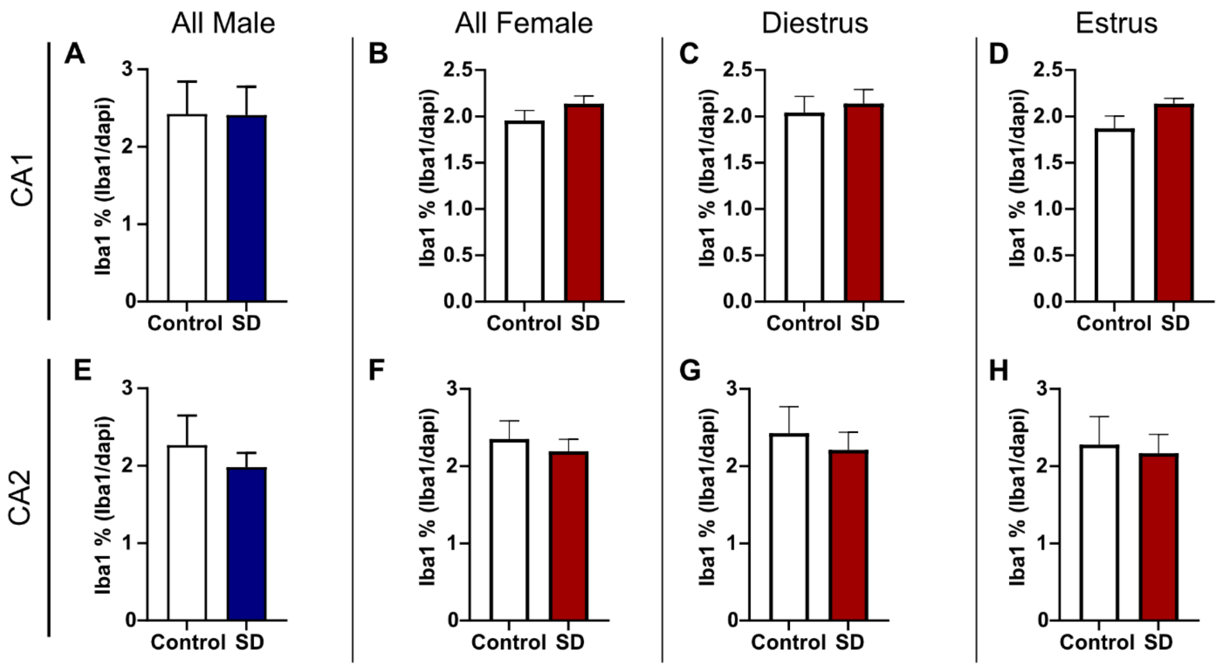


**Figure S3. Hippocampal pyramidal layer (PL) Iba1 expression is not affected by 120 h SD.** (A) There was no change in Iba1 expression in the hippocampal CA1 PL between male control and SD rats. (B-D) SD did not induce Iba1 expression among all females or based on estrous cycle stage. (E) No significant difference in Iba1 expression was found in the CA2 PL for male rats. (F-H) There was no difference in Iba1 expression between control and SD female rats regardless of estrous cycle stage. Bar graph data are presented as mean ± SEM.

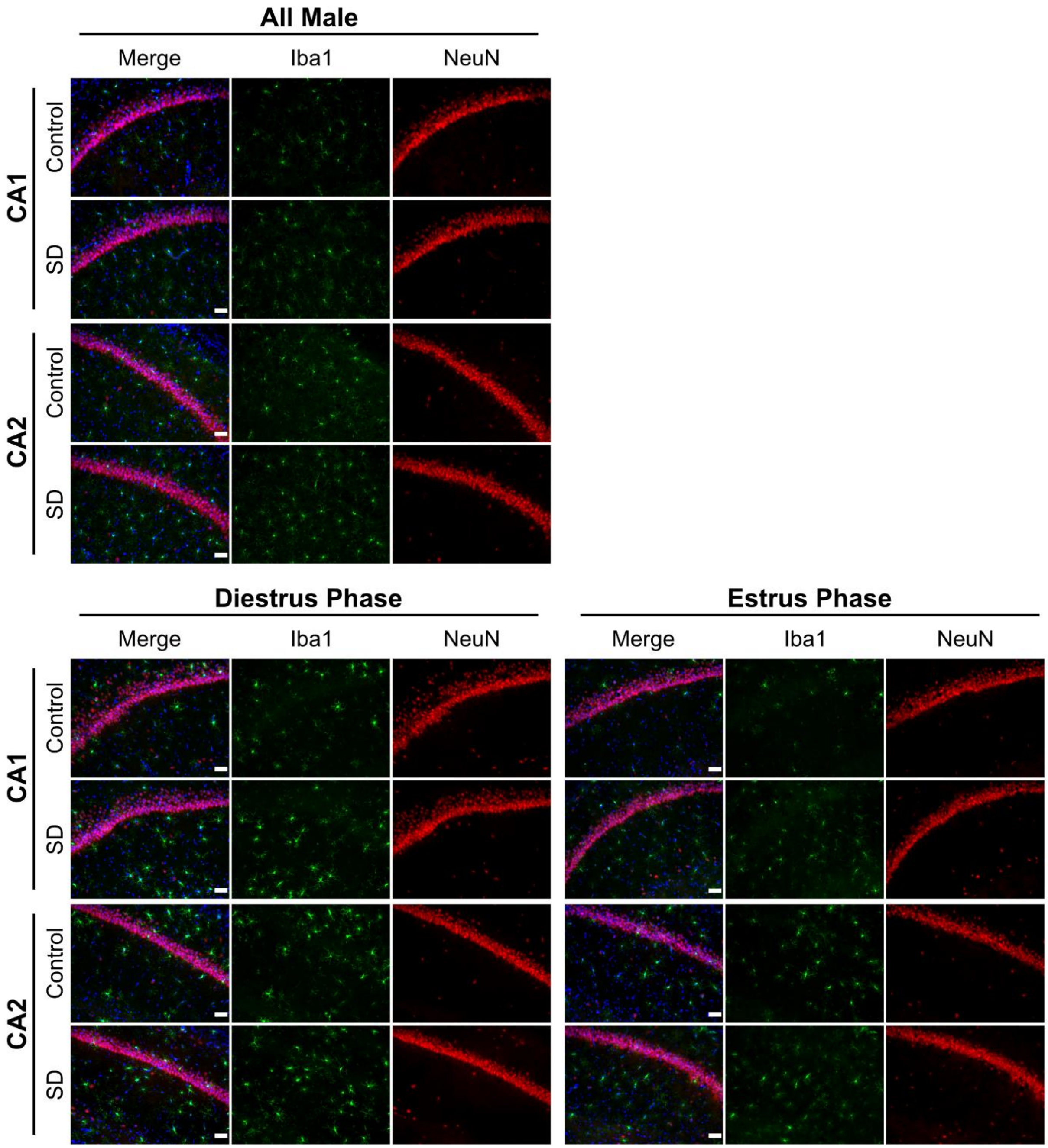


**Figure S4. Hippocampal Iba1 expression 120 h after SD.** Representative IHC images of Iba1 expression in the CA1 and CA2 as quantified in Figure 5 and Supplemental Figure 3. Scale bar = 50 µm.

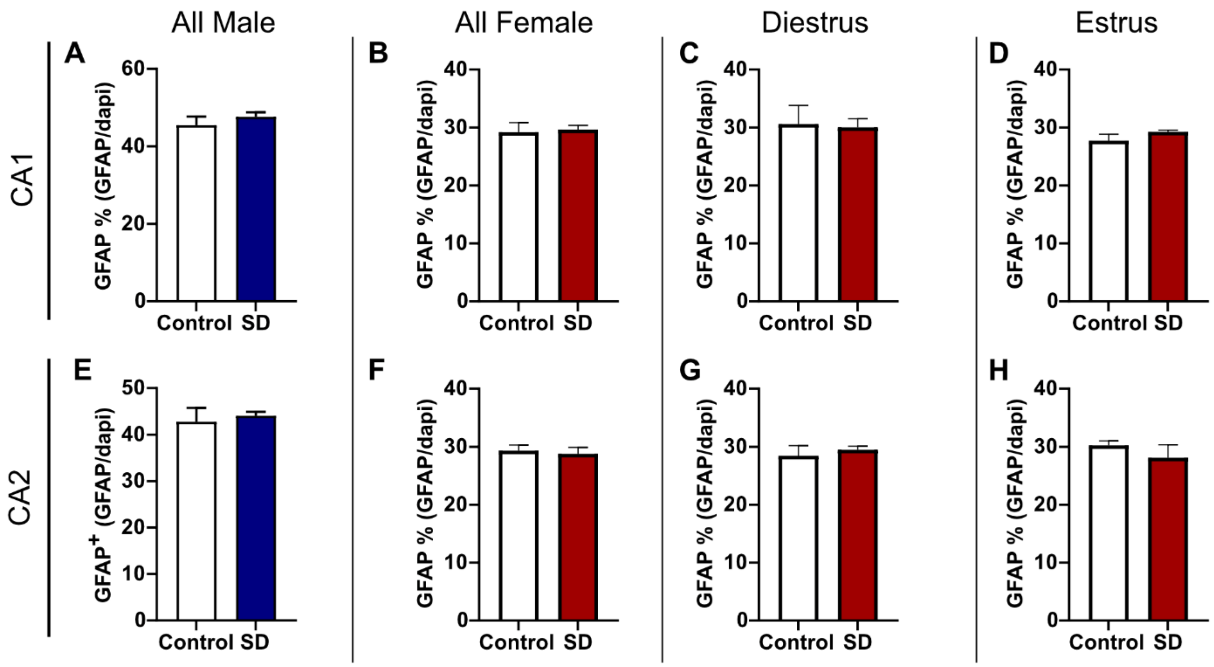


**Figure S5. GFAP expression in hippocampal SR is not affected by 120 h SD.** (A) There was no change in GFAP expression in the SR of the CA1 between male control and SD rats. (B-D) SD did not affect GFAP expression in the CA1 SR among all females or based on estrous cycle stage. (E) No significant difference in GFAP expression was found in the CA2 PL for male rats. (F-H) There was no difference in GFAP expression between control and SD female rats regardless of estrous cycle stage. Bar graph data are presented as mean ± SEM.

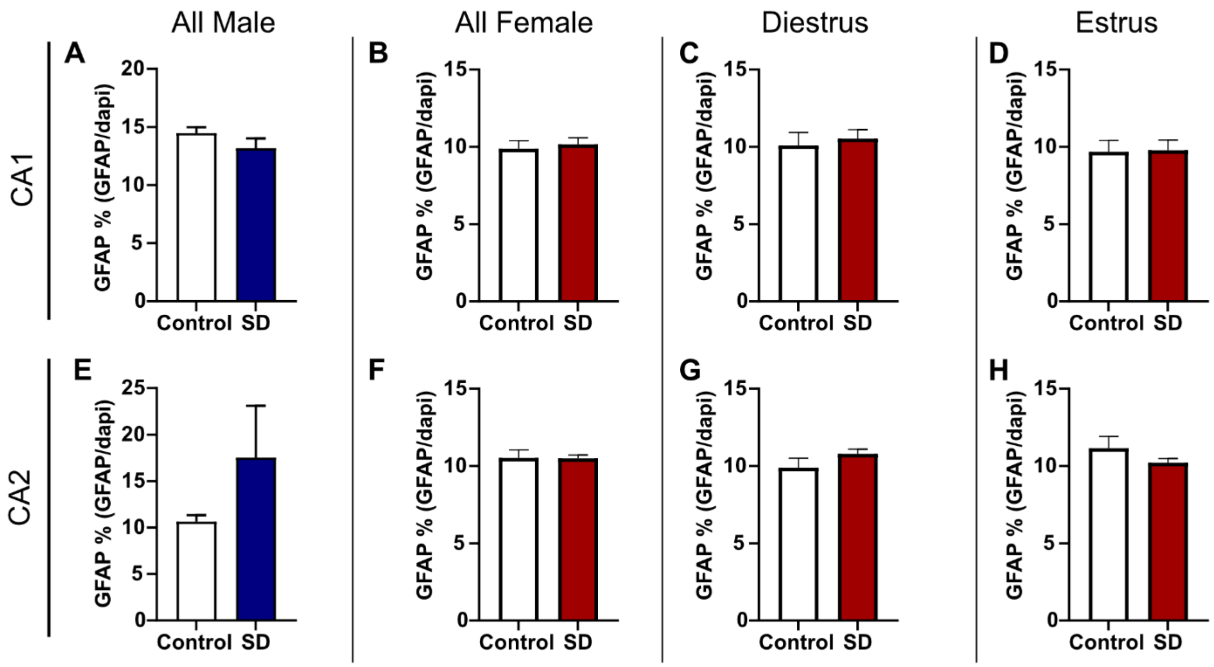


**Figure S6. Hippocampal PL GFAP expression is not affected by 120 h SD.** (A-B) SD did not change the total GFAP expression in the PL of the CA1 in male rats or female rats. (C-D) When estrous cycle was considered, there was no change in GFAP expression between control or SD female rats in the CA1 PL. (E) There was no significant difference in GFAP expression in the CA2 PL between control and SD male rats. (F-H) SD did not affect GFAP expression in the CA2 PL among all females or based on estrous cycle stage. Bar graph data are presented as mean ± SEM.

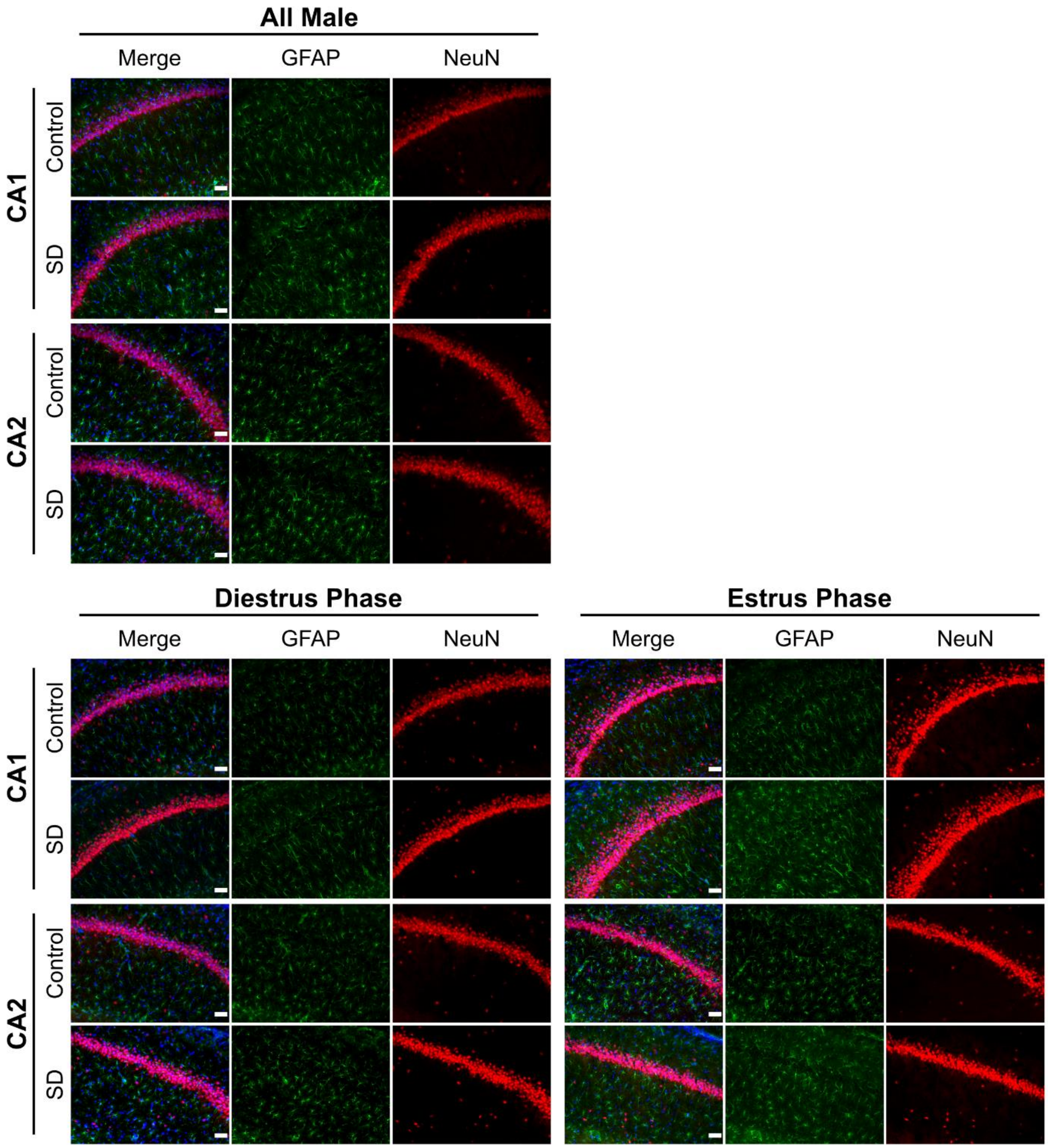


**Figure S7. Hippocampal GFAP expression 120 h after SD.** Representative IHC images of GFAP expression in the CA1 and CA2 as quantified in Supplemental Figure 5-6. Scale bar = 50 µm.

| Analysis | T | df | p-value | 95% CI of difference | Mean±SEM |
|---|---|---|---|---|---|
| ***Elevated Zero Maze- Male, All Subjects*** | | | | | |
| Distance Traveled | 4.629 | 19 | < 0.001 | 382.8 – 1015 | Control = 1781±90.74<br>SD = 2480±125.8 |
| Time in Open Arm(s) | 2.967 | 19 | 0.008 | 14.16 – 81.96 | Control = 121.6±11.82<br>SD = 169.7±9.987 |
| Head Dip Frequency | 3.957 | 19 | < 0.001 | 2.931 – 9.513 | Control = 11.67±1.170<br>SD = 17.89±0.9196 |
| Head Dip Time | 4.358 | 19 | < 0.001 | 6.011 – 17.12 | Control = 17.60±1.930<br>SD = 29.17±1.652 |
| ***Elevated Zero Maze- Female, All Subjects*** | | | | | |
| Distance Traveled | 1.053 | 34 | 0.300 | -169.0 – 532.5 | Control = 2749±118.4<br>SD = 2930±125.7 |
| Time in Open Arm(s) | 1.971 | 34 | 0.057 | -0.7369 – 47.80 | Control = 164.6±9.577<br>SD = 188.2±6.677 |
| Head Dip Frequency | 0.754 | 34 | 0.456 | -3.698 – 8.063 | Control = 34.05±2.098<br>SD = 36.24±1.964 |
| Head Dip Time | 1.107 | 34 | 0.276 | -4.014 – 13.63 | Control = 42.66±2.787<br>SD = 47.47±3.372 |
| ***Elevated Zero Maze- Diestrus Phase*** | | | | | |
| Distance Traveled | 0.886 | 25 | 0.384 | -254.8 – 639.6 | Control = 2715±152.1<br>SD = 2907±154.8 |
| Time in Open Arm(s) | 0.851 | 25 | 0.403 | -17.64 – 42.49 | Control = 172.3±12.07<br>SD = 184.7±7.758 |
| Head Dip Frequency | 0.123 | 25 | 0.903 | -6.823 – 7.691 | Control = 35.64±2.628<br>SD = 36.08±2.313 |
| Head Dip Time | 0.296 | 25 | 0.770 | -9.694 – 12.95 | Control = 45.42±3.418<br>SD = 47.05±4.360 |
| ***Elevated Zero Maze- Estrus Phase*** | | | | | |
| Distance Traveled | 0.623 | 7 | 0.553 | -453.1 – 777.5 | Control = 2843±161.6<br>SD = 3005±210.3 |
| Time in Open Arm(s) | 3.593 | 7 | 0.0088 | 19.21 – 93.14 | Control = 143.2±9.274<br>SD = 199.4±13.17 |
| Head Dip Frequency | 1.562 | 7 | 0.162 | -3.674 – 17.97 | Control = 29.6±2.379<br>SD = 36.75±4.211 |
| Head Dip Time | 3.527 | 7 | 0.0096 | 4.583 – 23.22 | Control = 34.94±2.489<br>SD = 48.84±3.133 |

**Table S1.** Statistical analyses for EZM behavioral data.

| Analysis | t | df | p-value | 95% CI of difference | Mean±SEM |
|---|---|---|---|---|---|
| ***Novel Object Recognition- Locomotor Activity*** | | | | | |
| Male, All Subjects | 3.113 | 19 | 0.006 | 1.041 – 5.314 | Control = 5.041±0.5848<br>SD = 8.218±0.8873 |
| Female, All Subjects | 0.749 | 38 | 0.458 | -1.821 – 0.8372 | Control = 10.09±0.4331<br>SD = 9.602±0.4932 |
| Diestrus Phase | 0.651 | 16 | 0.525 | -2.798 – 1.484 | Control = 10.14±0.8228<br>SD = 9.479±0.6135 |
| Estrus Phase | 0.340 | 20 | 0.737 | -2.275 – 1.637 | Control = 10.07±0.5240<br>SD = 9.751±0.8390 |

**Table S2.** Statistical analyses for NOR behavioral data.

| Analysis | F | df | p-value | 95% CI of difference | Means±SEM |
|---|---|---|---|---|---|
| ***Novel Object Recognition- Frequency of Object Exploration, Male All Subjects*** | | | | | |
| Main Effect Object | 0.570 | 1,13 | 0.464 | -6.435 – 3.102 | Familiar = 12.22±2.111<br>Novel = 13.89±0.444 |
| Main Effect Group | 1.568 | 1,13 | 0.233 | -6.964 – 1.853 | Control = 11.778±1.667<br>SD = 14.333±0.00 |
| Interaction: Object and Group | 0.570 | 1,13 | 0.464 | -12.87 – 6.203 | Familiar control = 10.111±1.837<br>Familiar SD = 14.333±1.687<br>Novel control = 13.444±1.676<br>Novel SD = 14.333±3.242 |
| ***Novel Object Recognition- Frequency of Object Exploration, Female All Subjects*** | | | | | |
| Main Effect Object | 1.879 | 1,27 | 0.181 | -4.697 – 0.9344 | Familiar = 11.71±1.104<br>Novel = 13.60±0.0404 |
| Main Effect Group | 0.701 | 1,27 | 0.410 | -3.948 – 1.660 | Control = 12.083±1.472<br>SD = 13.227±0.409 |
| Interaction: Object and Group | 0.600 | 1,27 | 0.445 | -7.758 – 3.505 | Familiar Control = 10.611±1.007<br>Familiar SD = 12.818±1.617<br>Novel Control = 13.556±1.027<br>Novel SD = 13.636±2.051 |
| ***Novel Object Recognition- Frequency of Object Exploration, Diestrus Phase*** | | | | | |
| Main Effect Object | 0.174 | 1,11 | 0.684 | -2.949 – 4.330 | Familiar = 12.67±1.333<br>Novel = 11.98±1.690 |
| Main Effect Group | 0.072 | 1,11 | 0.793 | -2.570 – 3.284 | Control = 12.50±1.167<br>SD = 12.143±1.857 |
| Interaction: Object and Group | 3.343 | 1,11 | 0.095 | -13.33 – 1.232 | Familiar Control = 11.333±1.308<br>Familiar SD = 14.0±2.0<br>Novel Control = 13.667±1.585<br>Novel SD = 10.286±0.778 |

| Analysis | F | df | p-value | 95% CI of difference | Means±SEM |
|---|---|---|---|---|---|
| ***Novel Object Recognition- Frequency of Object Exploration, Estrus Phase*** | | | | | |
| Main Effect Object | 10.06 | 1,14 | 0.0068 | -10.06 – -1.943 | Familiar = 10.50±0.250<br>Novel = 16.50±3.0 |
| Main Effect Group | 1.724 | 1,14 | 0.210 | -8.558 – 2.058 | Control = 11.875±1.625<br>SD = 15.125±4.375 |
| Interaction: Object and Group | 2.114 | 1,14 | 0.168 | -2.613 – 13.61 | Familiar Control = 10.250±1.388<br>Familiar SD = 10.750±2.780<br>Novel Control = 13.50±1.368<br>Novel SD = 19.50±4.291 |
| **Analysis** | **F** | **df** | **p-value** | **95% CI of difference** | **Means±SEM** |
| ***Novel Object Recognition- Object Exploration Time, Male All Subjects*** | | | | | |
| Main Effect Object | 0.767 | 1,13 | 0.397 | -3.567 – 1.509 | Familiar = 14.08±1.649<br>Novel = 15.11±1.287 |
| Main Effect Group | 0.032 | 1,13 | 0.861 | -4.740 – 4.015 | Control = 14.418±1.982<br>SD = 14.780±0.953 |
| Interaction: Object and Group | 6.243 | 1,13 | 0.027 | -10.95 – -0.7947 | Familiar Control = 12.436±1.347<br>Familiar SD = 15.733±2.161<br>Novel Control = 16.400±1.660<br>Novel SD = 13.827±1.214 |
| ***Novel Object Recognition- Object Exploration Time, Female All Subjects*** | | | | | |
| Main Effect Object | 1.335 | 1,27 | 0.258 | -6.143 – 1.717 | Familiar = 12.58±0.4391<br>Novel = 14.79±1.559 |
| Main Effect Group | 0.254 | 1,27 | 0.618 | -5.677 – 3.438 | Control = 13.128±0.108<br>SD = 14.247±2.105 |
| Interaction: Object and Group | 1.088 | 1,27 | 0.306 | -3.864 – 11.86 | Familiar Control = 13.020±2.271<br>Familiar SD = 12.142±1.875<br>Novel Control = 13.236±1.335<br>Novel SD = 16.353±2.452 |
| ***Novel Object Recognition- Object Exploration Time, Diestrus Phase*** | | | | | |
| Main Effect Object | 0.027 | 1,11 | 0.873 | -6.495 – 7.537 | Familiar = 13.71±0.4729<br>Novel = 13.19±0.3005 |
| Main Effect Group | 0.069 | 1,11 | 0.798 | -5.701 – 7.248 | Control = 13.833±0.347<br>SD = 13.060±0.174 |
| Interaction: Object and Group | .0029 | 1,11 | 0.958 | -13.69 – 14.38 | Familiar Control = 14.180±5.095<br>Familiar SD = 13.234±2.583<br>Novel Control = 13.487±2.120<br>Novel SD = 12.886±1.848 |
| ***Novel Object Recognition- Object Exploration Time, Estrus Phase*** | | | | | |
| Main Effect Object | 8.142 | 1,14 | 0.013 | -11.26 – -1.597 | Familiar = 11.34±1.105<br>Novel = 17.77±4.655 |
| Main Effect Group | 0.928 | 1,14 | 0.352 | -11.45 – 4.354 | Control = 12.775±0.335<br>SD = 16.325±6.095 |
| Interaction: Object and Group | 6.534 | 1,14 | 0.023 | 1.854 – 21.19 | Familiar Control = 12.440±2.442<br>Familiar SD = 10.230±2.640<br>Novel Control = 13.110±1.762<br>Novel SD = 22.420±4.888 |

**Table S3.** Statistical analyses for NOR behavioral data.

| Analysis | t | df | p-value | 95% CI of difference | Mean±SEM |
|---|---|---|---|---|---|
| ***Novel Object Preference- Novel Object Preference*** | | | | | |
| Male, All Subjects | 2.234 | 13 | 0.044 | -0.366 – -0.006 | Control = 0.1369±0.0561<br>SD = -0.0491±0.0572 |
| Female, All Subjects | 0.435 | 27 | 0.667 | -0.202 – 0.311 | Control = 0.07765±0.07319<br>SD = 0.1320±0.1064 |
| Diestrus Phase | 0.327 | 11 | 0.750 | -0.496 – 0.367 | Control = 0.07348±0.1407<br>SD = 0.009358±0.1356 |
| Estrus Phase | 1.561 | 14 | 0.141 | -0.0999 – 0.634 | Control = 0.07973±0.08899<br>SD = 0.3467±0.1236 |

**Table S4.** Statistical analyses for NOR NP behavioral data.

| Analysis | U | p-value | Median latency | | Mean±SEM |
|---|---|---|---|---|---|
| ***Passive Avoidance Task- Time to Cross*** | | | | | |
| Male, All Subjects | 17 | 0.007 | Control = 63.70, n=12<br>SD = 23.40, n=9 | | Control = 234.9±98.09<br>SD = 24.47±5.293 |
| **Analysis** | **T** | **df** | **p-value** | **95% CI of difference** | **Mean±SEM** |
| ***Passive Avoidance Task- Time to Cross*** | | | | | |
| Female, All Subjects | 3.189 | 38 | 0.003 | -122.2 – -27.29 | Control = 118.5±22.18<br>SD = 43.70±7.591 |
| Diestrus Phase | 1.513 | 16 | 0.150 | -148.2 – 24.76 | Control = 112.4±47.90<br>SD = 50.64±12.65 |
| Estrus Phase | 2.920 | 20 | 0.009 | -148.3 – -24.72 | Control = 121.7±24.05<br>SD = 35.22±6.583 |

**Table S5.** Statistical analyses for PAT behavioral data.

| Analysis | t | df | p-value | 95% CI of difference | Mean±SEM |
|---|---|---|---|---|---|
| ***Elevated Zero Maze- Diestrus vs Estrus phase controls*** | | | | | |
| Distance Traveled | 0.466 | 17 | 0.647 | -451.8 – 708.0 | Diestrus = 2715±152.1<br>Estrus = 2843±161.6 |
| Time in Open Arm(s) | 1.370 | 17 | 0.188 | -73.91 – 15.70 | Diestrus = 172.3±12.07<br>Estrus = 143.2±9.274 |
| Head Dip Frequency | 1.292 | 17 | 0.214 | -15.91 – 3.826 | Diestrus = 35.64±2.628<br>Estrus = 29.60±2.379 |
| Head Dip Time | 1.749 | 17 | 0.098 | -23.13 – 2.160 | Diestrus = 45.42±3.418<br>Estrus = 34.94±2.489 |
| ***Novel Object Recognition- Diestrus vs Estrus phase controls*** | | | | | |
| Locomotor Activity | 0.071 | 18 | 0.945 | -2.026 – 1.894 | Diestrus = 10.14±0.8228<br>Estrus = 10.07±0.5240 |
| Novel Object Preference | 0.039 | 16 | 0.969 | -0.333 – 0.3455 | Diestrus = 0.07348±0.1407<br>Estrus = 0.07973±0.08899 |
| ***Passive Avoidance Task- Diestrus vs Estrus phase controls*** | | | | | |
| Time to Cross | 0.196 | 18 | 0.847 | -90.92 – 109.6 | Diestrus = 112.4±47.90<br>Estrus = 121.7±24.05 |

| Analysis | U | p-value | Median latency | Mean±SEM |
|---|---|---|---|---|
| ***Elevated Zero Maze- Diestrus vs Estrus phase controls*** | | | | |
| Latency to Open Arm | 26 | 0.430 | Diestrus = 1.260, n=14<br>Estrus = 2.240, n=5 | Diestrus = 1.713±0.4688<br>Estrus = 2.568±0.9341 |

**Table S6.** Statistical analyses for diestrus vs estrus phase control behavioral data.

| Analysis | F | df | p-value | 95% CI of difference | Means±SEM |
|---|---|---|---|---|---|
| ***Novel Object Recognition- Frequency of Object Exploration, Diestrus vs Estrus controls*** | | | | | |
| Main Effect Object | 3.139 | 1,16 | 0.096 | -6.132 – 0.5486 | Familiar = 10.79±0.5417<br>Novel = 13.58±0.0833 |
| Main Effect Group | 0.161 | 1,16 | 0.693 | -2.675 – 3.925 | Control = 12.50±1.167<br>SD = 11.875±1.625 |
| Interaction: Object and Group | 0.084 | 1,16 | 0.775 | -5.764 – 7.597 | Familiar Control = 11.333±1.308<br>Familiar SD = 10.250±1.388<br>Novel Control = 13.667±1.585<br>Novel SD = 13.50±1.368 |
| ***Novel Object Recognition- Object Exploration Time, Diestrus vs Estrus controls*** | | | | | |
| Main Effect Object | 2.58x $10^{-5}$ | 1,16 | 0.996 | -4.856 – 4.880 | Familiar = 13.31±0.870<br>Novel = 13.30±0.1883 |
| Main Effect Group | 0.10 | 1,16 | 0.756 | -6.041 – 8.158 | Control = 13.833±0.347<br>SD = 12.775±0.335 |
| Interaction: Object and Group | 0.088 | 1,16 | 0.770 | -8.373 – 11.10 | Familiar Control = 14.180±5.095<br>Familiar SD = 12.440±2.442<br>Novel Control = 13.487±2.120<br>Novel SD = 13.110±1.762 |

**Table S7.** Statistical analyses for diestrus vs estrus phase control NOR data.

| Analysis | t | df | p-value | 95% CI of difference | Mean±SEM |
|---|---|---|---|---|---|
| ***Iba1 Expression- CA1, Stratum Radiatum*** | | | | | |
| Male, All Subjects | 3.014 | 9 | 0.015 | 0.5187 – 3.639 | Control = 8.586±0.4620<br>SD = 10.66±0.4969 |
| Female, All Subjects | 2.928 | 15 | 0.010 | 0.4820 – 3.062 | Control = 4.710±0.3261<br>SD = 6.482±0.4905 |
| Diestrus Phase | 1.287 | 7 | 0.239 | -1.075 – 3.643 | Control = 5.050±0.5231<br>SD = 6.334±0.7811 |
| Estrus Phase | 3.094 | 6 | 0.021 | 0.4803 – 4.115 | Control = 4.370±0.3813<br>SD = 6.668±0.6373 |
| ***Iba1 Expression- CA2, Stratum Radiatum*** | | | | | |
| Male, All Subjects | 3.319 | 8 | 0.011 | 0.8396 – 4.661 | Control = 8.068±0.4594<br>SD = 10.82±0.6894 |
| Female, All Subjects | 3.059 | 15 | 0.008 | 0.5161 – 2.887 | Control = 4.065±0.4365<br>SD = 5.767±0.3532 |
| Diestrus Phase | 1.423 | 7 | 0.198 | -0.9099 – 3.659 | Control = 4.188±0.8065<br>SD = 5.562±0.5794 |
| Estrus Phase | 3.379 | 6 | 0.015 | 0.5737 – 3.586 | Control = 3.943±0.4785<br>SD = 6.023±0.3873 |
| ***Iba1 Expression- CA1, Pyramidal Layer*** | | | | | |
| Male, All Subjects | 0.022 | 9 | 0.983 | -1.265 – 1.241 | Control = 2.423±0.4196<br>SD = 2.411±0.3657 |
| Female, All Subjects | 1.335 | 15 | 0.202 | -0.108 – 0.4687 | Control = 1.955±0.1070<br>SD = 2.136±0.085 |
| Diestrus Phase | 0.411 | 7 | 0.693 | -0.456 – 0.6479 | Control = 2.040±0.1757<br>SD = 2.136±0.1544 |
| Estrus Phase | | | | | |
| ***Iba1 Expression- CA2, Pyramidal Layer*** | | | | | |
| Male, All Subjects | 0.673 | 8 | 0.520 | -1.270 – 0.6960 | Control = 2.266±0.3832<br>SD = 1.979±0.1865 |
| Female, All Subjects | 0.589 | 15 | 0.565 | -0.747 – 0.4235 | Control = 2.354±0.2325<br>SD = 2.192±0.1563 |
| Diestrus Phase | 0.544 | 7 | 0.603 | -1.152 – 0.7213 | Control = 2.428±0.3422<br>SD = 2.212±0.2276 |
| Estrus Phase | 0.257 | 6 | 0.806 | -1.183 – 0.9578 | Control = 2.280±0.3626<br>SD = 2.168±0.2447 |

**Table S8.** Statistical analyses for IHC Iba1 data.

| Analysis | t | df | p-value | 95% CI of difference | Mean±SEM |
|---|---|---|---|---|---|
| ***GFAP Expression- CA1, Stratum Radiatum*** | | | | | |
| Male, All Subjects | 0.976 | 7 | 0.362 | -3.097 – 7.447 | Control = 45.47±2.197<br>SD = 47.64±1.164 |
| Diestrus Phase | 0.159 | 6 | 0.879 | -9.247 – 8.117 | Control = 30.60±3.218<br>SD = 30.03±1.496 |
| Estrus Phase | 1.299 | 6 | 0.242 | -1.337 – 4.362 | Control = 27.73±1.121<br>SD = 29.24±0.3164 |
| ***GFAP Expression- CA2, Stratum Radiatum*** | | | | | |
| Male, All Subjects | 0.511 | 8 | 0.623 | -4.631 – 7.268 | Control = 42.75±2.982<br>SD = 44.06±0.8382 |
| Diestrus Phase | 0.560 | 6 | 0.596 | -3.571 – 5.691 | Control = 28.42±1.788<br>SD = 29.48±0.6214 |
| Estrus Phase | 0.900 | 6 | 0.403 | -7.950 – 3.675 | Control = 30.23±0.7722<br>SD = 28.09±2.246 |
| ***GFAP Expression- CA1, Pyramidal Layer*** | | | | | |
| Male, All Subjects | 1.013 | 7 | 0.345 | -4.348 – 1.287 | Control = 14.47±0.5033<br>SD = 13.17±0.8501 |
| Female, All Subjects | 0.408 | 14 | 0.690 | -1.177 – 1.729 | Control = 9.878±0.5223<br>SD = 10.15±0.4315 |
| Diestrus Phase | 0.448 | 6 | 0.670 | -2.019 – 2.924 | Control = 10.09±0.8358<br>SD = 10.54±0.5670 |
| Estrus Phase | 0.100 | 6 | 0.923 | -2.338 – 2.538 | Control = 9.668±0.7382<br>SD = 9.768±0.6692 |
| ***GFAP Expression- CA2, Pyramidal Layer*** | | | | | |
| Male, All Subjects | 0.981 | 8 | 0.355 | -9.280 – 23.03 | Control = 10.64±0.7007<br>SD = 17.52±5.587 |
| Female, All Subjects | 0.045 | 14 | 0.965 | -1.217 – 1.167 | Control = 10.53±0.5117<br>SD = 10.50±0.2171 |
| Diestrus Phase | 1.303 | 6 | 0.240 | -0.7772 – 2.547 | Control = 9.903±0.6082<br>SD = 10.79±0.3025 |
| Estrus Phase | 1.146 | 6 | 0.295 | -2.931 – 1.061 | Control = 11.15±0.7690<br>SD = 10.22±0.2727 |

| Analysis | U | p-value | Median | Mean±SEM |
|---|---|---|---|---|
| ***GFAP Expression- CA1, Stratum Radiatum*** | | | | |
| Female, All Subjects | 20 | 0.234 | Control = 28.31, n=8<br>SD = 29.42, n=8 | Control = 29.16±1.668<br>SD = 29.64±0.7233 |
| ***GFAP Expression- CA2, Stratum Radiatum*** | | | | |
| Female, All Subjects | 28 | 0.721 | Control = 29.83, n=8<br>SD = 28.90, n=8 | Control = 29.32±0.9643<br>SD = 28.79±1.110 |

**Table S9.** Statistical analyses for IHC GFAP data.
